# Frequency-tagging captures distinct neural responses elicited by bilateral periodic thermonociceptive stimulation

**DOI:** 10.1101/2025.07.31.667970

**Authors:** Chiara Leu, Gabrielle Herbillon, Giulia Liberati, Valéry Legrain

**Author notes:** Corresponding author: Chiara Leu, Institute of Neuroscience, Université catholique de Louvain, Avenue Mounier 53, 1200 Bruxelles, Belgium.

## Abstract

Sustained periodic stimuli are known to elicit a periodic neural response (i.e. steady-state evoked potential) in the EEG frequency spectrum. These responses can easily be traced at their frequency of stimulation and corresponding harmonics using a frequency-tagging approach. To date, sustained periodic thermonociceptive stimuli have only been used on one extremity (e.g. right volar forearm) at a time. Extending this procedure to sustained stimulation applied concomitantly to distinct limbs would allow us to study the mechanisms of integration or competition between sensory signals from these different body locations. This study demonstrates that slow, sustained, sinusoidal thermonociceptive stimuli, bilaterally applied using two different stimulation frequencies (i.e. f_1_, f_2_, one on each forearm), elicit two distinct neural periodic responses at the respective frequency of stimulation and their harmonics. Additionally, we showed preliminary evidence for an interaction between the neural populations involved in the response to these stimuli, marked by neural activity at intermodulation frequencies (n* f_1_ ± m* f_2_). So far, this non-linear integration of sensory information has already been observed following visual and auditory stimuli but not yet following thermonociceptive stimuli.

**New and noteworthy:** This study demonstrates that sustained, slow, sinusoidal thermonociceptive stimulation applied simultaneously to both forearms at different frequencies elicits distinct neural responses at each stimulation frequency and its harmonics. Moreover, we provide preliminary evidence for interaction between the neural populations involved in the response to these stimuli during bilateral thermonociceptive stimulation. These findings extend frequency-tagging approaches in pain research and reveal potential non-linear sensory integration of distinct thermo-nociceptive inputs.

## 1. Introduction

Sustained periodic stimuli are known to elicit a periodic neural response at the frequency of stimulation and its harmonics (Regan, 1989). This frequency-tagging paradigm has been used for years to explore neural mechanisms underlying sensory perception domains such as auditory (e.g. (Geisler, 1960; Pantev et al., 1996)), vision (see Rossion et al. (2020) for a comprehensive review) or rhythm perception (Nozaradan et al., 2018; Nozaradan et al., 2011), but has also been extended to tactile (Moungou et al., 2016) and crucially, thermo-nociceptive stimuli (Colon et al., 2012a; Colon et al., 2012b; Mouraux et al., 2011).

Subsequent investigations demonstrated that an ultra-slow sustained periodic sinusoidal thermo-nociceptive stimulation applied at 0.2 Hz leads to improved signal-to-noise ratio (SNR) of the response to the nociceptive stimulation. Moreover, these slow steady-state evoked potentials (SSEPs) reflect predominantly the activation of slowly-adapting C-fibers (Colon et al., 2017). Being thus related to the *secondary* burning and lasting sensation of pain as well as being implicated in tissue inflammation (Campbell & Meyer, 1983), these SSEPs could potentially aid in the understanding of the role of C-fibers in pain perception (Iannetti et al., 2008). While this ultra-slow frequency-tagging paradigm has subsequently been used in a number of scalp EEG investigations (Leu et al., 2023; Leu et al., 2025; Leu et al., 2024; Mulders et al., 2020), which further deepened the understanding of these periodic brain responses, they have always been applied to one single body limb at a time (although sometimes concomitant with a stimulus of another sensory modality: e.g. Colon et al. (2014)). Yet, these applications do not tap into the full potential of this paradigm; given its high SNR generated, this type of slow sustained stimulus is highly suited to tease out otherwise relatively small differences in neural processing and integration of nociceptive stimuli. In particular, the concomitant application of these stimuli at different body sites, for instance bilaterally on both arms, would also allow us to assess characteristics of cognitive processing induced by e.g. spatial attention tasks (as demonstrated in other sensory modalities (Colon et al., 2015; Giabbiconi et al., 2007; Porcu et al., 2013; Saupe et al., 2009; Toffanin et al., 2009)).

On another note, concomitantly applied sensory periodic stimuli can elicit a phenomenon called intermodulation (*within* the same sensory modality) or crossmodulation (*between* sensory modalities), which is thought to reflect an interaction between the applied stimuli (i.e. f_1_ & f_2_), leading to the non-linear neural integration of the two input signals emerging at “intermodulation frequencies” in the EEG frequency spectrum (i.e. n* f_1_ ± m* f_2_) (Regan & Regan, 1988a; Regan & Regan, 1988b; Yang et al., 2016). While audiovisual crossmodulation at these specific frequencies was initially reported (Regan et al., 1995), subsequent studies have failed to replicate this effect using concomitant audiovisual (Giani et al., 2012), visual-tactile (Colon et al., 2015) or nociceptive-visual and nociceptive-tactile stimuli (Colon et al., 2014). Conversely, intermodulation by integration of different streams of visual input has been frequently achieved (reviewed in Chen et al. (2024); Gordon et al. (2019); Xu et al. (2022)). Using vibrotactile stimulation, intermodulation has also been found for concomitant stimulation on the same hand, but not when both hands were stimulated (Pang & Mueller, 2015). Yet, to the best of our knowledge, intermodulation has never been evidenced following concomitant sustained periodic nociceptive stimulation.

Thus, the main aim of this investigation was to demonstrate that the bilateral and concomitant application of ultra-slow sustained periodic sinusoidal thermo-nociceptive stimulation at two different frequencies (f_1_, f_2_) would elicit a periodic response in the EEG frequency spectrum for both applied stimulation frequencies. Upon confirmation of this effect, we aimed to further explore the possibility of intermodulation (i.e. non-linear integration of the concomitant stimuli), a response which would emerge at intermodulation frequencies, a combination of harmonics of f_1_ and f_2_ (i.e. n* f_1_ ± m* f_2_).

## 2. Methods

This experiment was pre-registered on the Open Science Framework (OSF, https://doi.org/10.17605/OSF.IO/KN5QE) prior to the beginning of data collection.

### 2.1. Participants

Twenty healthy participants (between 18 and 35 years old) were recruited via an established website and received compensation of 15€ for their participation in the experiment (duration approximately 1.5 h). The data of two participants had to be removed from the analysis due to technical problems with the EEG recordings and were subsequently also removed from the behavioral data analysis. The remaining participants (n = 18) were 24.2 ± 3.5 (mean ± std. dev.) years old (range: 20-35) and 8 participants were female. Only right-handed participants were included in the investigation to avoid any potential confound of handedness of the participants in our results.

Exclusion criteria included regular use of psychotropic medication, intake of 1st level analgesics (i.e. paracetamol and nonsteroidal anti-inflammatory drugs) within 12 hours before the experiment), as well as any severe neurological diseases, psychiatric disorders, recent trauma or skin disease affecting upper limbs. All participants gave written informed consent prior to the beginning of the assessment and were informed that they could withdraw from the experiment at any given time. All procedures were performed in accordance with the relevant guidelines and regulations. The local research ethics committee approved all experimental procedures (Commission d’Ethique hospitalo-facultaire Saint-Luc UCLouvain, B403201316436).

The software G*Power was used to conduct a power analysis. We aimed to reach a power of 0.8 using a standard 0.05 alpha error probability. Based on data of Colon et al. (2017), we expected a relatively large effect size (d=1.15) for the detection of a periodic response (tested against zero, one-sided) at the frequency of stimulation, resulting in a suggestion of 10 participants. To account for a potentially smaller effect size given the concomitant bilateral stimulation (as opposed to the commonly used unilateral 0.2 Hz stimulation), to match investigations in other domains which observed responses at the intermodulation frequencies (Boremanse et al., 2013; Giani et al., 2012), and to account for potential dropouts, we increased the sample size to 20 participants.

### 2.2. Stimuli and materials

Two thermal cutaneous stimulators (TCS II with T03 probe, QST.Lab, Strasbourg, France) were used to deliver sustained periodic thermonociceptive stimuli on the volar forearms of the participants. The probes were set with 15 micro-Peltier elements (each ∼7.7 mm2, arranged in a 30-mm diameter circle), whose temperature can vary at rates of up to 300 °C/sec. The temperature of stimulation varied between 35 °C and 50 °C in a sustained periodic sinusoidal manner. Two frequencies of stimulation were applied for 60 sec per trial, one on each arm of the participant: Stimulation at frequency 1 (f_1_) was delivered at 0.25 Hz (i.e. containing 15 cycles of 4 sec), while stimulation at frequency 2 (f_2_) was delivered at 0.333 Hz (i.e. containing 20 cycles of 3 sec) (Figure 1). The forearm on which f_1_ and f_2_ were applied was counterbalanced during the experiment, so that the left and right arm received the same number of stimuli with both stimulation frequencies across the experiment. Inter-trial-intervals were variable and self-paced by the experimenter to allow participants to provide ratings of intensity and painfulness. The thermodes were displaced after each trial to avoid habituation or sensitization. Stimulation frequencies were determined based on previous studies demonstrating that ultra-slow sustained periodic thermonociceptive stimuli have a superior SNR in frequency-tagging paradigms compared to faster paced thermonociceptive stimulation (Colon et al., 2017). Additionally, the frequencies were chosen so that their harmonics would not overlap in the frequency spectrum. Lastly, we aimed to have stimulation frequencies that are not too different from each other to avoid delivering significantly more stimulation cycles on one arm than on the other during the same trial (i.e. avoiding confounds due to e.g. difference in terms of temporal summation).

**Figure 1.**
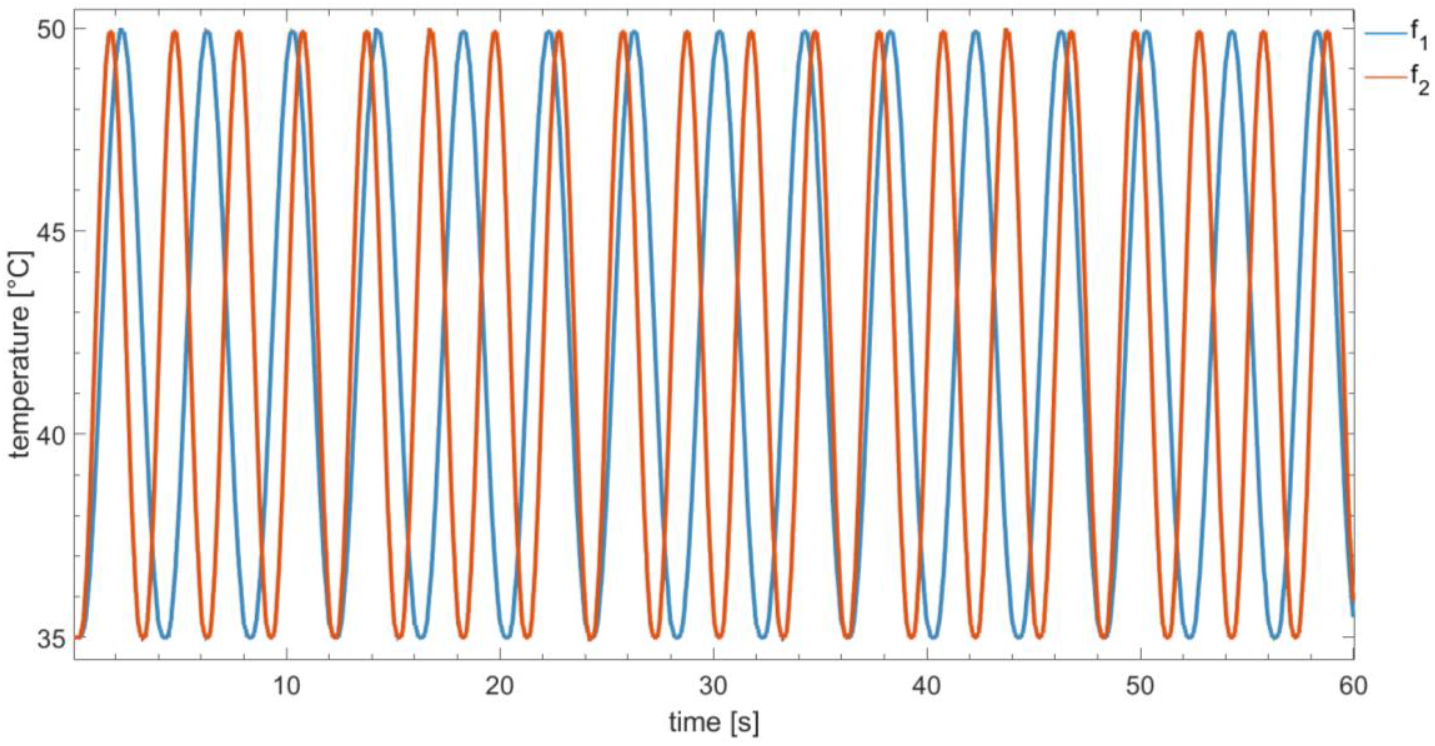
Illustration of the bilateral and concomitant stimulation. Sustained periodic sinusoidal thermonociceptive stimuli were delivered at the frequencies f_1_ (0.25 Hz) and f_2_ (0.333 Hz), using a thermal cutaneous stimulator (TCS II, QST.Lab, Strasbourg, France).

### 2.3. Procedure

Participants were seated comfortably in a chair in front of a table, on which both their forearms were resting with the volar surface upwards. They were instructed to move as little as possible and keep their gaze constant to avoid interference with the EEG signal acquisition during the thermonociceptive stimulation. Additionally, they were encouraged to focus their attention on the bilaterally applied stimulation during the trials. In total, 30 trials of the bilateral thermonociceptive stimulation were delivered, split into 6 blocks of 5 stimulation trials. The arm on which f_1_ and f_2_ were applied was randomized and counterbalanced across blocks, so that each arm received an equal amount of stimulation using f_1_ and f_2_.

### 2.4. Behavioral data

Participants were asked to give a global rating of the intensity of their perception of the bilateral thermonociceptive stimulation on a numerical rating scale (NRS) ranging from 0 (“no perception”) to 10 (“most intense perception imaginable”) after each trial. Participants were encouraged to give a spontaneous rating, reflecting their perception as adequately as possible. Additionally, they were asked after each trial to evaluate whether they perceived the stimulation as painful, answering either with “yes” or “no”. Finally, after each block (i.e. 5 trials), participants were asked whether they perceived the left and right stimulation differently from each other.

### 4.5. EEG data

EEG signals were recorded using an elastic electrode-cap with 64 active, pre-amplified Ag-AgCl electrodes arranged according to the international 10-10 system; signals were digitalized at 1024 Hz and stored for offline analyses (ActiveTwo, BioSemi, Netherlands). To maintain a clear signal, the direct-current offset was limited to 30 mV. All electrodes were re-referenced offline to the average electrode activity.

The EEG recordings were preprocessed using the Letswave7 (www.letswave.org) toolbox and analyzed using custom scripts in MATLAB 2022a (The MathWorks Inc., Natick, Massachusetts).

#### 2.5.1. Preprocessing

To process EEG data, we employed a frequency-tagging analysis approach (Regan, 1966; Regan, 1989) to analyze the periodic response induced by the slow sustained periodic stimuli, which allows for an objective differentiation between oscillatory activity related to the applied stimuli and other unrelated ongoing activity (Colon et al., 2012b). The frequency-tagging method is based on the notion that a periodic stimulus elicits a periodic activity which can be identified as periodic responses at the frequency of stimulation in the recorded EEG signals (Colon et al., 2012a; Mouraux et al., 2011). This approach has been frequently used in our lab, leading to a standardized analysis approach (Colon et al., 2017; Leu et al., 2023; Leu et al., 2025; Leu et al., 2024; Mulders et al., 2020).

The obtained EEG signal was first filtered using a fourth-order Butterworth band-pass filter between 0.05 and 100 Hz, then a Notch filter with a 2 Hz width was applied at 50 and 100 Hz. The signal was segmented into epochs of the length of stimulation (60s), relative to the onset of the stimuli. Due to their placement on the EEG cap, electrodes Iz, P9 and P10 are often very noisy and were therefore removed from the data set to improve signal quality. To remove potential muscular artifacts (i.e., from eye movements), an Independent Component Analysis (Fast ICA algorithm) (Hyvarinen & Oja, 2000) was applied, and any trial containing amplitudes larger than ± 500 μV was removed. On average, ∼ 3 ± 2 independent components supposed to reflect noise signals were removed per participant. Less than 1% of trials were removed overall due to large artifacts that could not be removed using the ICA. Noisy channels were interpolated using their 3 neighboring electrodes. The remaining signal was re-referenced to the average of the electrode set and epochs were concatenated to a length of 120 sec, to improve the signal resolution in the frequency domain (frequency resolution = 1/epoch duration, i.e., 1/120 sec = 0.00833 Hz). The epochs were then averaged across participants. To analyze the signal in the frequency domain, a discrete Fourier transform (FFT) (Frigo & Johnson, 1998) was used. Finally, residual noise was partially removed by subtracting the average amplitude of the signal measured at 2–8 neighboring frequencies (thereby excluding the bins immediately adjacent to the bin of interest), at each electrode and at each frequency bin (Mouraux et al., 2011).

#### 2.5.2. Aggregation of harmonics

The neural response to periodic stimulation is known to be distributed across harmonics in the frequency spectrum, and these harmonics should be aggregated (i.e., summed up) for an adequate reflection of the neural response to the stimulation (Retter et al., 2021). While there are many different approaches on how to define the ideal number of harmonics to sum, we chose to find the harmonic at which the aggregated signal has the highest signal-to-noise ratio (SNR) in the frequency spectrum, using baseline-corrected group- and channel-averaged data for f_1_ and f_2_. The SNR was calculated by dividing the signal at a given frequency bin *x* by the mean amount of noise at the 8 surrounding frequency bins (*x / baseline noise*, excluding immediately adjacent bins). This method differs slightly from previous investigations from our lab, where the harmonics were either not considered (Colon et al., 2017; Mulders et al., 2020) or all harmonics in the frequency spectrum were aggregated, regardless of their contribution to the SNR (Leu et al., 2023; Leu et al., 2024). Here, we adopted an intermediate approach that retains relevant harmonic components of the brain’s response while limiting the inclusion of noise thereby providing a more balanced and reliable estimate of the signal

To aggregate the harmonics, a custom MATLAB script was used to sum up the amplitude at each harmonic of each frequency of stimulation consecutively and assess the new SNR at each step (evaluated against 8 frequency bins on each side). Based on this evaluation, the highest SNR was identified at 9.25 Hz for f_1_ and 7.333 Hz for f_2_. Similar to other thermonociceptive SSEP investigations, the first few harmonics carried most of the periodic neural response (Colon et al., 2017; Leu et al., 2023; Leu et al., 2025; Leu et al., 2024; Mulders et al., 2020).

A multi-sensor cluster-based Wilcoxon signed-rank test (clusters built using up to 4 electrode neighbors, 1000 permutations, p<0.05, right-tailed) was used to objectively assess the electrodes with the largest responses (i.e. amplitude) at the aggregated harmonics for f_1_ and f_2_ separately.

#### 2.5.3. Intermodulation responses

To assess intermodulation responses, the baseline-corrected signal was grand averaged across participants, and z-scores were calculated at each frequency bin by calculating the difference between the amplitude of the frequency bin and the averaged amplitude of the 8 frequency bins on each side (16 in total), excluding the immediately adjacent bins, and divided by the standard deviation of these 8 bins (following the formula: *x – baseline noise / std dev*_*baseline noise*_).

Significant z-scores at intermodulation frequencies (i.e. n* f_1_ ± m* f_2_,where *n* and *m* are integers) were identified in the group-averaged EEG spectrum up to 50 Hz at each channel *(p<0*.*001)*. For each channel and participant, the baseline subtracted amplitudes at the significant intermodulation frequencies of that channel were extracted and (after excluding direct harmonics of f_1_ and f_2_) aggregated to form the “intermodulation response” at that channel. Thus, for each channel, a different number of intermodulation harmonics was aggregated to create an accurate reflection of the neural response to the bilaterally applied stimuli across the scalp.

#### 2.5.4. Assessment of the number of trials necessary for a good SNR

Finally, a larger number of trials were used in this experiment than in previous frequency-tagging investigations from our lab. To explore how many trials would suffice for a good SNR at the frequency of stimulation and harmonics of f_1_ and f_2_ (excluding the intermodulation response), we ran the analysis for either the first 6, 12, 18 and 24 trials of each participant and assessed the SNR of the baseline-subtracted group- and channel-averaged signal at the subsequently aggregated harmonics in the frequency spectrum

### 2.5 Statistical analyses

Statistical analyses were carried out using R Statistical Software (Version 4.3.1, R Core Team 2023) and MATLAB (2022a). The significance level was set at *p≤*0.001 for the identification of significant intermodulation responses, consequently z-scores ≥ 3.1 were considered as significant (one-sided, signal > noise). For other statistical tests the significance level was set to *p* ≤ 0.05.

Linear mixed models (LMM) including *subject* as random effect (accounting for intercept variations between individuals) were used to assess differences in intensity ratings across the stimulation blocks. A generalized linear mixed effects model with *subject* as random intercept and a binomial error distribution was used to assess the binary ratings of pain perception, testing overall block effects with a Wald χ^2^ test. All pairwise comparisons versus block 1 were obtained from model-estimated marginal means with Holm-adjusted p-values. A Wilcoxon two-sample paired signed-rank test was used to evaluate whether the aggregated amplitudes elicited by stimulation with f_1_ and f_2_ differed from each other.

## 3. Results

### 3.1. Behavioral data

On average, the stimuli were perceived as painful in 35 % of the trials (Figure 2A). 5 participants never perceived the stimulation as painful, while for most other participants the number of trials that were perceived as painful varied between the stimulation blocks. The mixed-effects logistic regression did not indicate a significant main effect of stimulation block on whether stimuli were perceived as painful (χ^2^(5) = 9.260, p = 0.099). Relative to block 1, odds of a perceiving a trial as painful were not significantly different in block 2 (odds ratio (OR) = 1.83, 95% CI [0.59, 5.74], p = 0.34), block 3 (OR = 1.83, 95% CI [0.59, 5.74], p = 0.34), or block 6 (OR = 2.45, 95% CI [0.78, 7.67], p = 0.13). Odds were marginally higher in block 4 (OR = 2.97, 95% CI [0.95, 9.31], p = 0.056) and significantly higher in block 5 (OR = 3.27, 95% CI [1.04, 10.27], p = 0.038). All participants indicated that they perceived the stimuli at a similar intensity on both arms. Overall, the stimuli were evaluated at an intensity of 5.33 ± 2.28 (mean ± std. dev.). A significant main effect of stimulation block was found in the LMM (F(5, 517) = 3.722, p = 0.003, η_p_^2^ = 0.03). Post hoc comparisons of model estimated marginal means (EMM) showed that compared to block 1, mean intensity ratings did not differ significantly in block 2 (EMM = 0.26, 95% CI [–0.18, 0.71], p = 0.13) or block 3 (EMM = 0.36, 95% CI [–0.08, 0.81], p = 0.073). However, intensity perception ratings were significantly higher in block 4 (EMM = 0.55, 95% CI [0.10, 1.00], p =0.005), block 5 (EMM = 0.58, 95% CI [0.14, 1.03], p = 0.003), and block 6 (EMM = 0.60, 95% CI [0.15, 1.05], p = 0.003).

**Figure 2.**
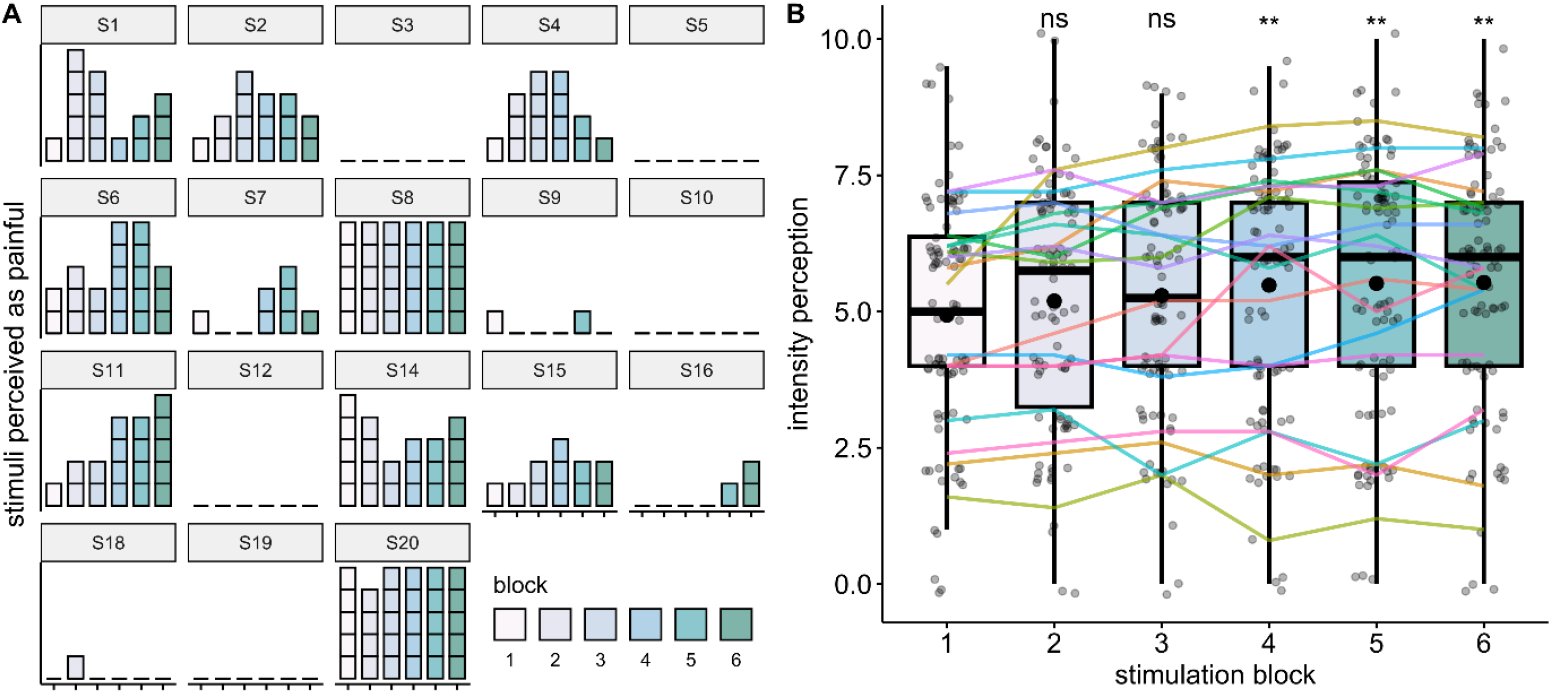
Ratings of perceived painfulness and intensity of the heat pain stimuli. **a)** Illustration of the number of trials that were perceived as painful for each participant and each stimulation block. **b)** Ratings of perceived stimulation intensity in each block. Horizontal lines indicate the evolution of each participants’ average rating per block, while the mean for each block is indicated using round black dots. Individual data points (i.e. data for each trial and each participant) are jittered around the boxplots in grey. Results of the post-hoc comparison of the model estimated marignal means comparing each block to block 1 are indicated: _ns_ p>0.05, ** p<0.01.

### 3.2. EEG data

Stimuli applied with frequency f_1_ and f_2_ elicited clear periodic responses at their respective frequency of stimulation and their harmonics (Figure 3). The first few harmonics were the largest for both stimulation frequencies, but they were distributed slightly differently across the frequency spectrum, leading to a different number of harmonics that were aggregated to achieve the highest SNR for the response to each stimulation intensity.

**Figure 3.**
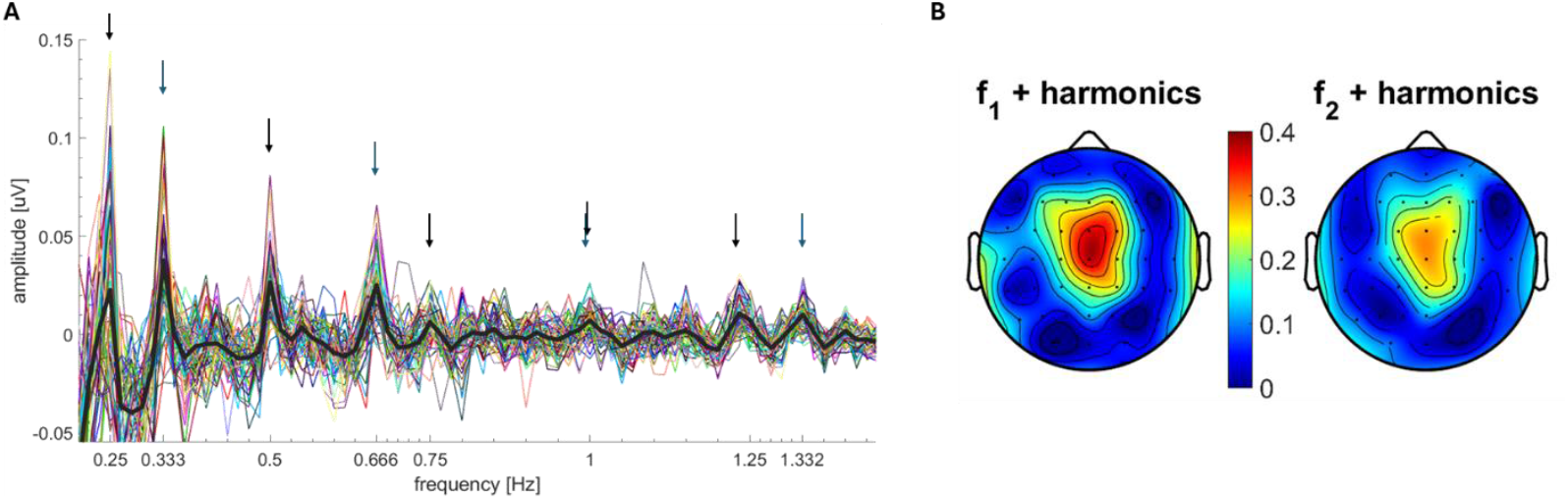
Neural response recorded following concomitant and bilaterally applied sustained periodic thermonociceptive stimulation on the volar forearms. **A)** Steady-state evoked potentials elicited by stimuli at frequency f_1_ (.25) and f_2_ (.333) in the EEG frequency spectrum. Group averaged data are indicated for each channel (colored) and the pooled channels (black). **B)** Topographical plots represent the response at the aggregated harmonics for f_1_ and f_2_, based on the number of aggregated harmonics with the highest SNR in the frequency spectrum estimated from group- and channel-averaged data for f_1_ and f_2_.

The aggregated harmonics demonstrated activity mainly distributed around the scalp vertex (f_1_: W(Cz) = 3.40; f_2_: W(Cz) = 3.57) at both frequencies of stimulation (Figure 3B) as well as at temporal channels in f_1_ (W(TP7) = 3.70; W(T8) = 3.48). The aggregated amplitudes at the scalp vertex elicited by stimulation with f_1_ were slightly larger than the ones elicited by f_2_ (W= 138, p = 0.021, n = 18, r = 0.54).

As we used a rather large number of trials in this experiment, we also explored how many trials would suffice to achieve a good SNR, having in mind that future studies will likely employ multiple experimental conditions while using bilateral stimuli. This sub-analysis on the first 6, 12, 18 and 24 trials of the experiment revealed that at least 12 trials are required to obtain a comparatively good SNR for stimuli at f_1_. Interestingly, stimulation frequency f_2_ seems to have overall a higher absolute SNR if more than 18 trials are included. The SNR of 12 trials is similar to f_1_ with the same number of trials (Figure 4).

**Figure 4.**
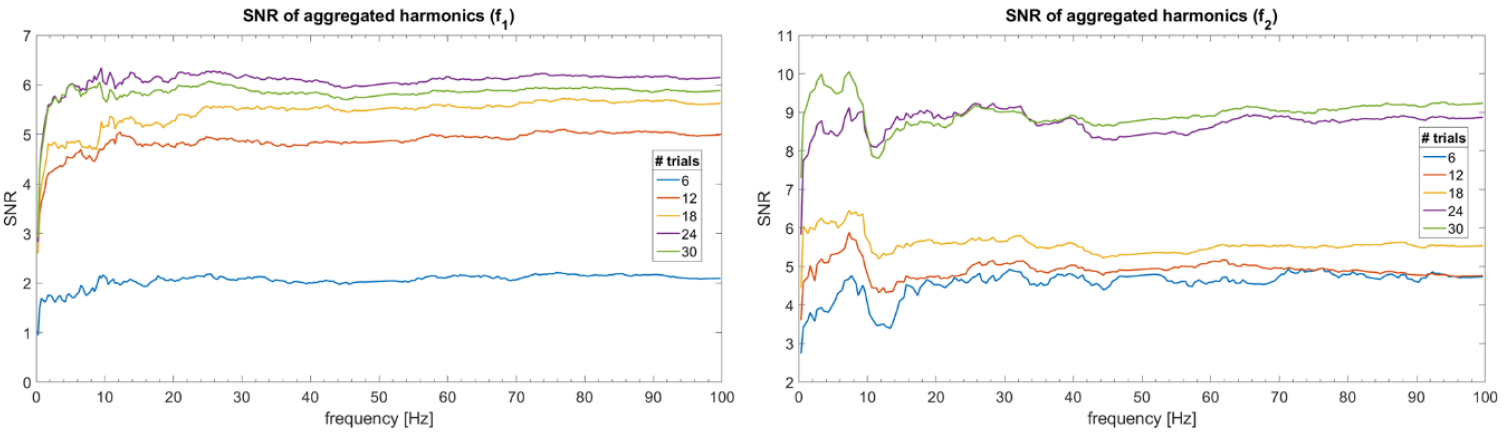
SNR development across the frequency spectrum for different numbers of trials. For each frequency of stimulation, the number of trials averaged before the frequency transformation was gradually decreased to assess its impact on the SNR across the frequency spectrum. The aim was to find the lowest number of trials still achieving a comparatively high SNR.

Significant intermodulation frequencies were identified based on their z-scores, separately for each channel. The amplitudes at these frequencies were aggregated to quantify the intermodulation response across the scalp. The largest intermodulation responses were found at parietal-central channels (Figure 5A). Most participants exhibited an intermodulation response (signal > noise) at almost all channels, with multiple harmonics being present (see example of channel Pz in Figure 5B and 5C).

**Figure 5.**
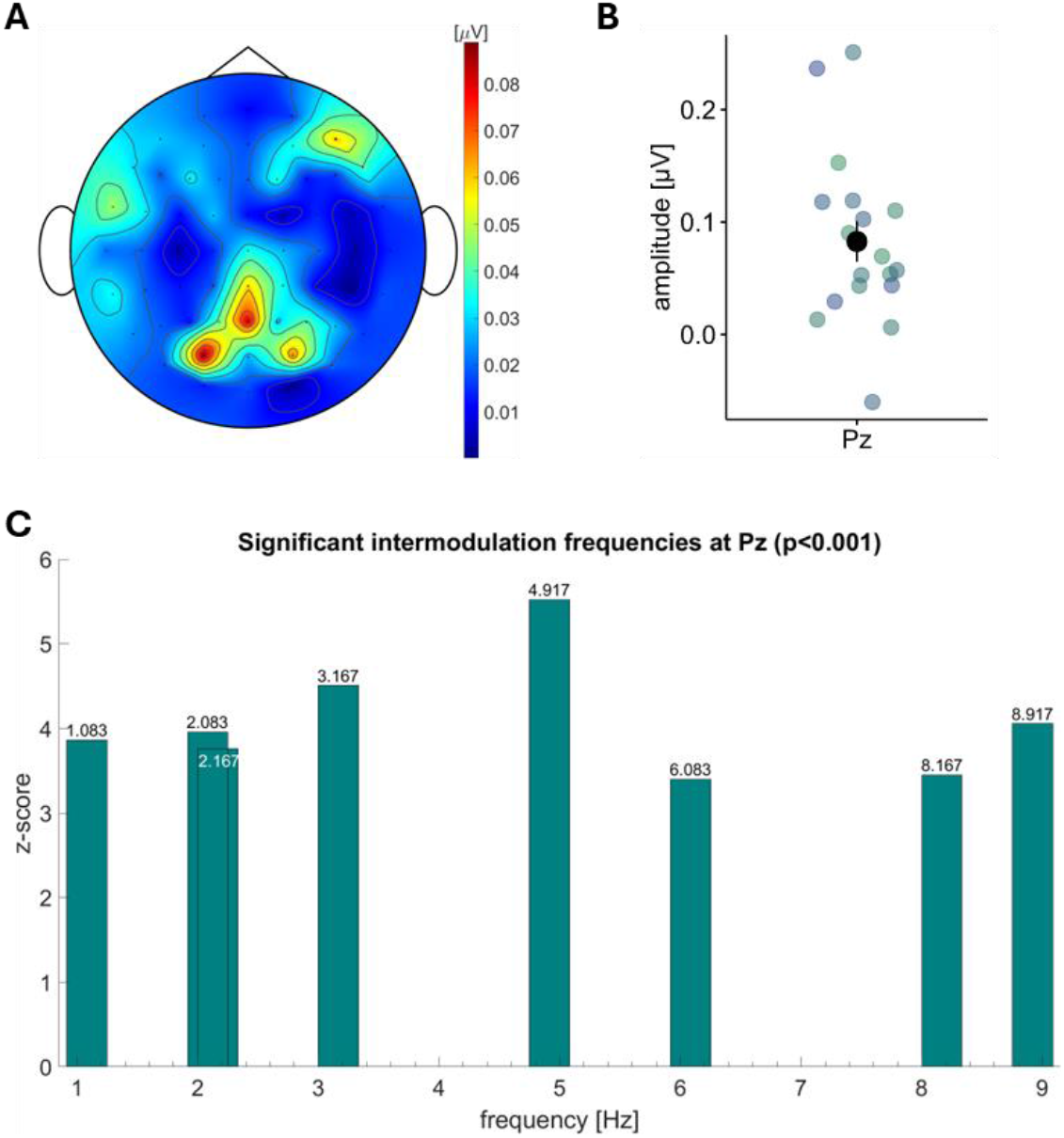
Neural activity at intermodulation frequencies. **A**) Topographic illustration of the group-level amplitude of the aggregated significant intermodulation frequencies for each channel. **B)** Individual participant aggregated baseline-subtracted amplitudes at channel Pz. **C)** Group-level z-scored responses at electrode Pz exceeding the threshold of z ≥ 3.1 (p < 0.001) at intermodulation frequencies up to 10 Hz. The specific frequencies of the significant z-scores are indicated in the plot.

## 4. Discussion

### 4.1. Bilateral frequency-tagging of sustained periodic thermonociceptive stimuli

This proof-of-concept study aimed to demonstrate that bilateral and concomitant application of ultra-slow sustained periodic sinusoidal thermonociceptive stimulation at frequencies f_1_ and f_2_ elicits periodic neural responses (i.e. SSEPs) at each respective frequency of stimulation (and its harmonics), which can objectively identified using a frequency-tagging approach. Additionally, we investigated whether the simultaneous activation of potentially similar neuronal populations by the bilateral stimulation leads to a non-linear integration of the nociceptive input, which would be present at intermodulation frequencies (i.e. n* f_1_ ± m* f_2_).

Both stimuli elicited clear SSEPs in the frequency spectrum at the frequency of stimulation and their harmonics. The largest periodic response to these concomitantly applied ultra-slow stimuli was found at the scalp vertex for both stimuli, consistent with other thermonociceptive SSEP investigations (Colon et al., 2017; Leu et al., 2023; Leu et al., 2025; Leu et al., 2024; Mulders et al., 2020), and likely reflecting activity of the ACC as well as left and right operculo-insular cortices (Mouraux et al., 2011). The obtained frequency spectra further parallel the previous thermonociceptive SSEP investigations, as the first few harmonics carried most of the periodic neural response. While the magnitude of the aggregated response differed slightly between f_1_ and f_2_, this is likely due to the relatively large difference in the number of harmonics that were aggregated in each response.

Previous studies comparing time-locked event-related potentials (ERPs) elicited by bilateral vs. unilateral application of brief thermonociceptive laser stimuli reported inconsistent results. In one experiment, ERP magnitudes elicited by bilateral stimulation decreased compared to unilateral stimulation (Northon et al., 2019), while they increased (particularly when the hands were close to the body) in another (Northon et al., 2022). One reason for the variability in these results may be that since ERPs are obtained by averaging EEG signals in the time domain and therefore reflect superposition of the different responses indiscriminately, this does not allow to understand the individual contribution of the ascending signals generated by each limb stimulation. Using a SSEP paradigm with two distinct stimulation frequencies could address this, allowing for clearer differentiation of input from each stimulated arm. Future studies could directly compare unilateral and bilateral stimulation to better understand how the brain processes concomitant versus single streams of thermonociceptive stimulation.

Given the relatively large number of trials in this experiment, we examined how reducing trial count would affect the SNR at the frequencies of stimulation and their harmonics. This is particularly relevant for future studies, where multiple conditions make high trial numbers impractical due to participant discomfort and long sessions. Our analysis showed that fewer than 30 trials were sufficient, with at least 12 trials providing good SNR at both frequencies, consistent with similar unilateral thermonociceptive stimulation studies.

Another reason to reduce the number of trials is the observed increase in stimulus intensity ratings over the course of the experiment, with significantly higher ratings in blocks 4–6 compared to block 1. This increase is likely related to the phenomenon of peripheral sensitization. Although the location of the thermode was shifted after each stimulus, given the total number of 30 stimuli, repeated stimulation of nearby areas with the same stimulation was unavoidable, potentially leading to spatial summation (Nielsen and Arendt-Nielsen, 1997). This observed sensitization emphasizes the need to limit the number of stimuli in future SSEP studies using ultra-slow sustained periodic thermonociceptive stimulation. Curiously, the number of trials perceived as painful is about half of the percentage of trials reported as painful in previous investigations (Leu et al., 2023; Leu et al., 2024), despite similar levels of intensity perception. The repetitive and predictive nature of the applied stimuli might have contributed to a decrease in the level of *perceived bodily threat* of these stimuli. In the previously mentioned thermonociceptive SSEP investigations, variations in conditions were always present, therefore leading to more variations in perception and potentially reducing the predictability of the stimuli.

A limitation of this experiment is that the chosen frequencies of stimulation are challenging to assess in the frequency spectrum due to the characteristics of frequency f_2_ (i.e. frequency of 0.333 Hz). The resolution in the frequency domain is limited by the duration of the trial; thus, a stimulation frequency with a higher precision than the frequency resolution is likely to succumb to rounding errors, as harmonics may fall into adjacent frequency bins rather than precisely where expected. This makes it more difficult to identify the exact responses to the applied stimuli, particularly when trying to aggregate harmonic responses. An improvement for future investigations would be to use a similarly slow frequency of stimulation with a more convenient value, e.g. 0.4 Hz, so that harmonics align more precisely with frequency bins and can be analyzed more reliably.

### 4.2. Intermodulation

With the successful implementation of the bilateral frequency-tagging using thermonociceptive stimuli, we set out to identify significant responses at intermodulation frequencies, which would illustrate the non-linear neural integration of the concomitantly applied stimuli. Since the neural activity elicited by these stimuli was most prominent at the scalp vertex, we expected that their activity might *overlap* and create an interaction, leading to the neural integration of the input of the two applied stimuli. This hypothesis was also supported by the perception of the participants, which – while not necessarily being perceived *as one* – also did not differ between the stimuli. This could have also been facilitated by the use of a global perception rating, rather than asking participants for a separate rating for each arm. Using a standard z-score threshold we found a large number of significant z-scored frequency bins across the frequency spectrum. Thus, to be more selective and forego issues of multiple comparison, we increased the significance threshold to a more conservative p < 0.001. Still, most channels showed multiple significant intermodulation harmonics, predominantly at higher harmonic combinations (i.e. higher than 1 Hz).

The main difference to previous attempts at investigating the non-linear integration of nociceptive stimuli might lie in the improvement in SNR (Colon et al., 2017). In the first attempts to elicit SSEPs through nociceptive stimulation, relatively fast (> 3 Hz) thermonociceptive (Colon et al., 2014; Mouraux et al., 2011) and low-intensity intraepidermal electrical (Colon et al., 2012b) stimuli were used, predominantly activating Aδ-fibers with maximal responses over fronto-central regions. Comparing these responses to concurrently applied tactile and visual stimulation showed that each modality elicited activity in a distinct subgroup of neuronal populations (Colon et al., 2012a), but that selectively attending either thermonociceptive or tactile stimulation did not alter the magnitude of the SSEP responses (Colon et al., 2014). One caveat of these studies was the relatively low SNR in the response to the high-frequency sustained periodic thermonociceptive stimuli. Given the improved SNR using ultra-slow stimuli (Colon et al., 2017), it could be interesting to revisit previous attempts at investigating potential cross-modal interactions between activation of the nociceptive system and other sensory stimuli and thereby improve our understanding of how a salient and threatening stimulus modulates and integrates with other sensory information.

The phenomenon of intermodulation following concomitant and periodic sensory input is not new. In the visual domain, the usefulness of neural integration of two different stimuli has elegantly been demonstrated by periodically presenting half-faces at different frequencies, during which the brain tries to integrate the information to a complete face (Boremanse et al., 2013, 2014). Comparatively, it is less clear what the responses recorded at intermodulation frequencies reflect in the context of bilateral thermonociceptive stimulation. Usually, intermodulation frequencies can be found only few harmonic combinations away from their “base” frequency (e.g. 1*f_1_+1*f_2_). This was not the case in this investigation, where intermodulation frequencies were identified predominantly at higher harmonic combinations, which raises the question of what processes these responses reflect. The fact that no intermodulation responses were found at early harmonic combinations in the z-scored signal shows that these responses were relatively small. This could have been partially caused by the timing of the two stimuli, which were never truly simultaneous (i.e. the applied heat cycles never reached their maximum exactly at the same time, though close to each other) and peaked sometimes relatively far from each other, which could have hampered a potential cleaner neural integration output. In an exploratory analysis, we isolated stimulation cycles that peaked almost simultaneously (see Figure 1, first cycle of the stimulation) and analyzed only this portion of the signal. The results were encouraging, showing a clearly identifiable peak at the first intermodulation frequency with the expected topography. However, we cannot be certain that the correct f_2_ frequency was identified, as aligning cycles to f_1_ disrupted the regularity of f_2_ in the time domain, potentially splitting it into two frequencies after transformation.

Thus, these higher harmonic responses may reflect interhemispheric coherence as the brain integrates sensory input from each arm, but given their small size at unusually high harmonics, they warrant cautious interpretation. Although this study cannot resolve the issue conclusively, our findings provide a foundation for future research on how the human brain integrates concomitant thermonociceptive activation, either alone or in combination with other sensory modalities.

## 5. Conclusion

This investigation successfully demonstrated that sustained periodic thermonociceptive stimuli applied concomitantly to both forearms at a frequency f_1_ and f_2_ elicit a periodic neural response at both frequencies of stimulation, easily identifiable using frequency-tagging. This paves the way for extending the frequency-tagging paradigm to a bilateral application of (thermo-)nociceptive stimuli, a promising paradigm to better understand the neural representation and integration of concomitant nociceptive stimuli. We further demonstrated the presence of periodic responses at intermodulation frequencies, which evidences the non-linear integration of the concomitantly applied thermonociceptive stimulation. Thus, using slow sustained periodic thermonociceptive stimulation could be beneficial not only to better understand nociceptive processing by itself, but also its cognitive modulation and multisensory integration in relation to other sensory stimuli.

## Acknowledgments

We wish to thank Francesca Barbero and members of the Pain Research Lab at UCLouvain for their helpful feedback and suggestions. The preprint version of this article is available athttps://doi.org/10.1101/2025.07.31.667970.

## Data availability

Behavioral and EEG data for this study are publicly available at https://doi.org/10.17605/OSF.IO/6QTUR.

## Author Contributions

Conceived and designed research: CL, GH, VL

Analyzed data: CL

Performed experiments: CL, GH

Interpreted results of experiments: CL

Prepared figures: CL

Drafted manuscript: CL

Edited and revised the manuscript: CL, GH, GL, VL

Approved final version of the manuscript: CL, GH, GL, VL

